# Mycolic acid and cholesterol induced transcriptomic adaptations in wild type and Mce complex ATPase subunit deleted strain of *M. tuberculosis* H37Rv

**DOI:** 10.1101/2024.05.13.593965

**Authors:** Mohammad Asadur Rahman, Valerio Izzi, Lee Riley, Rajaram Venkatesan

**Affiliations:** Faculty of Biochemistry and Molecular Medicine, University of Oulu, Oulu, Finland; Faculty of Medicine, BioIM Unit, University of Oulu, Oulu, Finland; School of Public Health, University of California, Berkeley, USA

**Keywords:** Mycobacterium tuberculosis, Mce-ABC transporters, Mkl, knock-out, RNA-seq, mycolic acid, cholesterol

## Abstract

During infection and latency, *Mycobacterium tuberculosis* (Mtb) primarily relies on lipids as its main energy source. The Mtb mammalian cell entry (Mce) complexes function as lipid transporters with Mkl as the ATPase subunit for substrate translocation. Cholesterol and mycolic acid are essential to the infectiousness and pathogenicity of Mtb. Here, we have investigated the transcriptomic responses of wild type and *Δmkl* strains of Mtb to cholesterol and mycolic acid through RNAseq analysis. The number of differentially expressed genes was always higher when grown on cholesterol than on mycolic acid. In most cases, *Δmkl* showed the opposite gene expression pattern compared to the wild type. Among the four *mce* operons, significant changes in expression patterns were observed mainly in the genes of *mce3* and *mce4* operons. The upregulation of *mce3* operon and the downregulation of *mce4* operon in the presence of mycolic acid and cholesterol is intriguing. The transcriptome profile of genes involved in mycolic acid synthesis did not show significant change to either mycolic acid or cholesterol. However, interestingly, both mycolic acid and cholesterol induce upregulation of methyl citrate cycle and dormancy related genes indicating their significance during Mtb persistence and provides insights on Mtb’s adaptive strategies under stress through transcriptional remodelling.

## 1. Introduction

*Mycobacterium tuberculosis* (Mtb) is the pathogen responsible for more than a million deaths every year and one-fourth of the human population is latently infected with TB [1]. Mtb could persist lifelong in hosts as latent infections where the bacilli continuously replicate retaining low metabolic activity and reactivation could happen anytime due to the host’s weakened immune system [2, 3]. The prolonged persistence of Mtb in the human host is due to its very distinctively thick and protective cell wall, mostly composed of lipids [4]. Mycolic acid is a prominent lipid found in the cell wall of mycobacteria to maintain the cellular architecture and impermeability [5, 6] and is important in its virulence, immunological processes, and antibiotic resistance [7-9]. Mtb tends to conserve energy and resources by recycling or modifying imported mycolic acids or synthesizing it from host fatty acids [10-12]. Cholesterol is essential to the infectiousness and pathogenicity of Mtb [13, 14]. Mtb uses fatty acids and cholesterol to produce metabolic intermediates or fuel metabolism or to utilize for biosynthesis directly [15, 16]. During infection, Mtb utilizes cholesterol as a carbon source and is important for its persistence [17, 18]. As a kind of nutrition recycling, Mtb may be able to receive fatty acids from the host cell membranes throughout an infection [19]. Alternatively, Mtb may also be able to obtain the lipids from the dying bacteria [10]. The influence of host lipids, particularly cholesterol, on the dynamics of infection and the precise functional significance of its metabolism by Mtb, remains inadequately elucidated [20]. There is also a limited understanding of mycolic acid recycling/reuse as a carbon source. First-line anti-tuberculous medicines target their biosynthesis pathway and cell envelope, though some of these pathway targets are unexplored [7].

Mtb genome has around 250 lipid metabolism-related genes, but their specific roles in metabolic pathways remain to be explored. Mtb has four homologous operons *mce1-4* involved in the transport of lipids across the cell wall to utilize them as sources of carbon and energy. The Mce1 complex encoded by the *mce1* operon is involved in fatty acid/mycolic acid transport. Mce operon functions as a ABC transporter and Mkl (also known as MceG) (Rv0655) is the ATPase to energize all four Mce complexes for substrate translocation [21-23].

Understanding the transcriptome is essential for figuring out how the genome functions, revealing the molecular components and molecular details of cells [24]. RNAseq (RNA sequencing) is a robust method for higher resolution transcriptomic measurement and is advantageous over quantitative reverse transcription-PCR or microarrays [25, 26]. Recent studies indicate that the lipid-rich environment significantly influences the Mtb transcriptome to a greater extent [26, 27]. Instead of using normal growth media containing dextrose or glycerol, fatty acid conditions could give a more realistic view of transcriptomic changes during infection [25]. Several studies have been done to understand the effect of fatty acid and cholesterol at the transcriptomic level. Rodriguez et al. have used a mixture of fatty acids containing oleic, palmitic and stearic acids [28], whereas in another study cholesterol, along with a mixture of fatty acids was used as the sole carbon source [25]. Recently, Pawelczyk and coworkers used only cholesterol as the sole carbon source and compared the transcriptomic changes in respect to glycerol [26].

In our study, we investigate the changes in transcriptomic levels by utilizing cholesterol or mycolic acid as the sole carbon source. Furthermore, we also studied the potential impact of *mkl*, the ATPase subunit of the lipid importing Mce1 complex on overall transcriptional changes by comparing the gene expression profile of *Δmkl* strain grown in glycerol or cholesterol or mycolic acid as the carbon source. The response of Mtb towards cholesterol and mycolic acid is not well understood. Though mycolic acid is the major and unique constituent of the cell wall and can be recycled, the effect of mycolic acid has not been measured yet/lacking. To our knowledge, this is the first study to understand the effect of mycolic acid and the *mce1* operon controlling ATPase gene, *mkl* in the global transcriptional responses and global metabolic adaptations in Mtb.

## 2. Materials and methods

### 2.1 Bacterial Strains and Growth Conditions

*Mtb* strain H37Rv and derivative strains (*Δmkl*), *M. smegmatis* were grown in either Sauton’s medium with 2% glycerol or modified Sauton’s medium (another carbon source) or Middlebrook 7H9 broth containing 10% ADC/OADC, 0.2% glycerol and 0.05% Tween-80 or Middlebrook 7H10/7H11 agar containing 10% OADC, 0.5% glycerol. Hygromycin at 50 μg ml^-1^ was used where appropriate. Frozen stocks were made from a single culture and used throughout the experiments. Mtb H37Rv related experiments were conducted in BSL3 and BSL2 laboratories at the University of California, Berkeley, USA.

### 2.2 Construction of an Mtb *mkl* mutant strain

An in-frame *mkl* knockout was made in Mtb H37Rv strain (a hygromycin cassette) using a temperature-sensitive phage-mediated recombineering method with slight modification [29, 30]. In brief, 500 bp 5’ and 3’ flanking regions of *mkl* were cloned at the MCS regions on either side of the *loxP-hygR-loxP* region of pMSG360. This vector was linearized with BsrBI and DraI restriction enzymes and the linear recombineering substrate was gel purified. Recombinase was expressed in EL350/phAE87 *E. coli* strain by incubating in a water bath at 42 ºC for 20 min and the linearized substrate was electroporated into it and plated on LB-agar plates supplemented with 150 µg ml^-1^ hygromycin and incubated for 2 days at 30 ºC. Hygromycin-resistant colonies were pooled and sub-cultured overnight and phagemid DNA was isolated from them by phenol-chloroform extraction method. Phagemid DNA was then transformed into *M. smegmatis* mc^2^155 by electroporation, mixed with top agar (0.6% w/v agarose in ddH_2_O, 2 mM CaCl_2_) on 7H10 plate and incubated at 30 ºC for 2 days. Plaques were collected from *M. smegmatis* lawns and amplified and high-titer phage was produced. Phages were collected using MP buffer (50 mM Tris pH 7.6, 150 mM NaCl, 10 mM MgCl_2_, 2 mM CaCl_2_) and filter sterilized using 0.2 µM filter syringe and stored at 4 ºC. Mtb was transduced with phage at 37 ºC for 4-6 h and plated on 7H11 plate with 100 µgml^-1^ hygromycin and incubated at 37 ºC for 4 weeks. Colonies that appeared were cultured again in hygromycin containing 7H9 media for genomic DNA extraction and samples were analyzed by sequencing.

### 2.3 Total RNA extraction

Total RNA was extracted from 30 ml of 7-day-old cultures of Mtb (wild type, *Δmkl*), grown in minimal media with glycerol or cholesterol or mycolic acid as a carbon source, according to a standard Trizol RNA extraction protocol (Invitrogen) in BSL3 lab. Then In-column DNase I Treatment and RNA cleanup was done using the Monarch RNA Cleanup Kit (NEB). Then total RNA was sent to QB3-Berkeley Genomics facility at the University of California, Berkeley, USA for quality checking, library preparation and sequencing.

### 2.4 RNAseq analysis

The Galaxy platform (https://usegalaxy.eu) was used for all sequences’ pre-processing and alignment [31, 32], using *ad-hoc* General Feature Format (GFF) and Gene Transfer Format (GTF) files downloaded from Mycobrowser (2021 release, https://mycobrowser.epfl.ch/releases). FastQc (https://www.bioinformatics.babraham.ac.uk/projects/fastqc/) was used to check the quality of the sequenced RNA data, followed by TrimGalore (https://www.bioinformatics.babraham.ac.uk/projects/trim_galore/) to trim the adapter sequences, hierarchical indexing for spliced alignment of transcripts 2 (Hisat2) for alignment [33, 34] and FeatureCount, a read summarization program for counting reads generated by RNAseq experiments [35].

Differential gene expression (DGE) analysis was performed in R with DESeq2 [36], and principal component analysis (PCA) was employed to detect outliers and reveal potential inter- and intra-treatment clusters. The Database for Annotation, Visualization and Integrated Discovery (DAVID) (https://david.ncifcrf.gov) [37] was used to annotate the results, based on gene ontology (GO, www.geneontology.org) lists for biological processes (BP), molecular function (MF), and cellular component (CC). Pathway enrichment analyses were based on the Kyoto Encyclopaedia of Genes and Genomes (KEGG) database (www.genome.jp). Results with a p-value of ≤ 0.05 were considered as significant. Further functional gene annotation was performed using the categories provided by Mycobrowser.

## 3. Results

### 3.1 Differential gene expression (DGE) analysis

To investigate the impact of lipids, specifically cholesterol and mycolic acid, on Mtb transcriptional dynamics we compared the gene expression patterns of Mtb grown with cholesterol or mycolic acid as the sole carbon source against glycerol. In parallel, another comparison was made in gene expression levels between wild type and *Δmkl* strains when using glycerol, cholesterol or mycolic acid as the sole carbon source, and DGEs were classified into the different functional categories as listed in mycobrowser [38]. PCA analysis and heatmaps show that carbon sources and biological replicates are the only factors affecting gene expression in our results. PCA analysis reveals, also, a clear separation between the samples grown with glycerol and the remaining samples, while cholesterol and mycolic acid samples form separate clusters. Notably, there are significant differences among cholesterol and mycolic acid wild type and mutant replicates themselves, unlike glycerol samples showing minimal changes according to the heat map. The consistency of biological replicates suggests stable samples during sequencing, indicating that the results are unaffected by technical issues and increases the reliability of the results (Fig. 1). In this study, DGEs were selected based on log2 fold change (FC) of ≥ 1 or ≤−1 (fold change of 2) and adjusted p-value/false recovery rate of ≤ 0.05, to identify statistically significant DGEs.

**Fig. 1.**
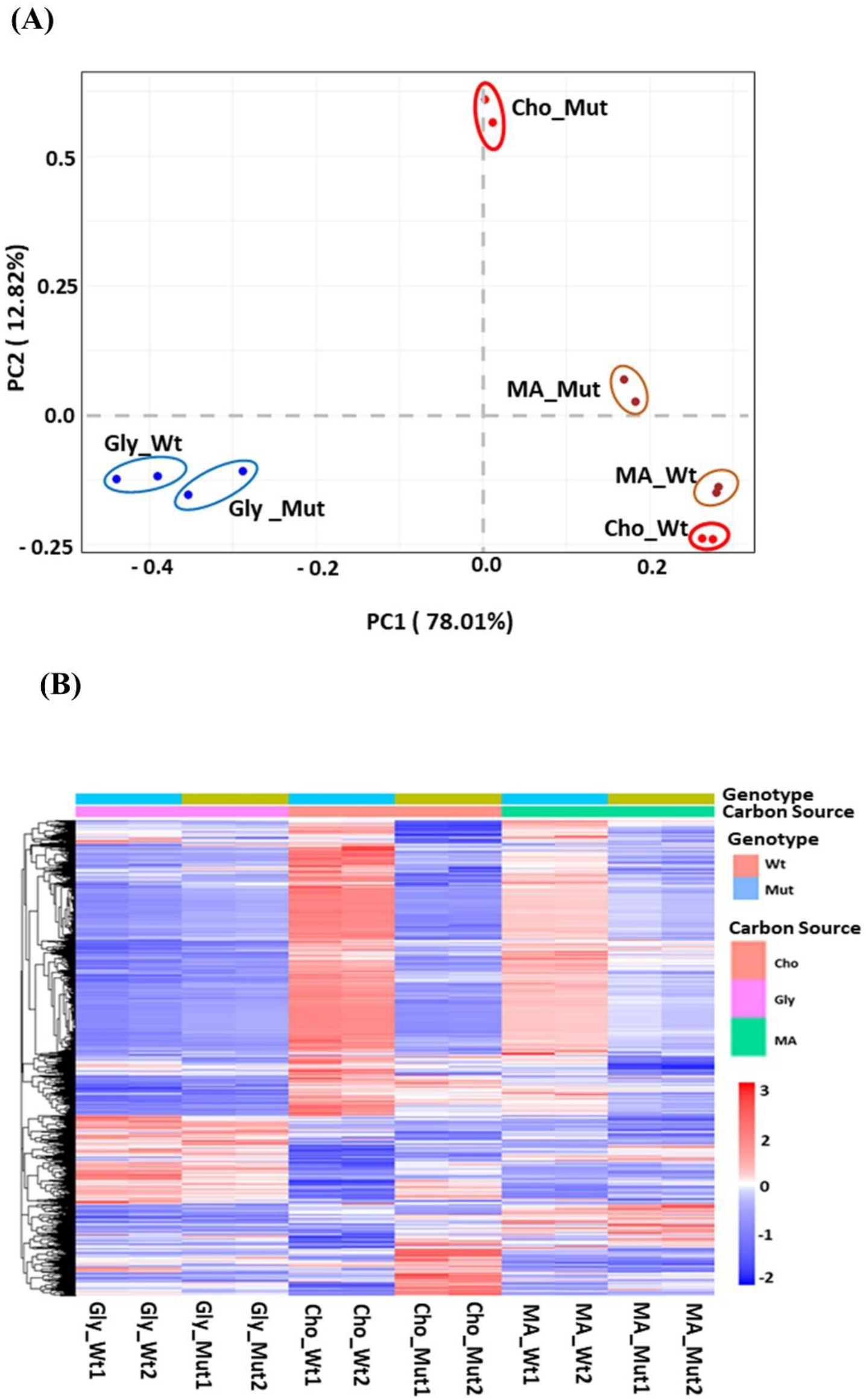
PCA plot and clustering heatmap. A). PCA plot analysis of the experimental conditions with their biological replicates. For the PCA plot, the data was rlog transformed. B). The clustering-heatmap. For the clustering heatmap, normalized counts were rescaled between −2 and 3. Clustering was based on Pearson correlation.

The DGE analysis reveals that both cholesterol and mycolic acid show drastic changes in wild type Mtb H37Rv. However, the number of genes exhibiting differential expression in the presence of cholesterol is higher (1408 genes; 722 upregulated and 686 downregulated) when compared to that in the presence of mycolic acid (938 genes; 478 upregulated and 460 downregulated). Interestingly, among the differentially expressed genes in the cholesterol and mycolic acid conditions, 642 genes are present in both conditions (Fig. 2). In both cases, the functional category analysis reveals that a significant number of differentially expressed genes belong to the categories of conserved hypothetical proteins/unknown, intermediary metabolism and respiration and cell wall and cell processes (Table 1). Furthermore, among the differentially expressed genes in the presence of cholesterol, 109 genes were related to lipid metabolism, with 53 genes upregulated and 56 genes downregulated, whereas in the presence of mycolic acid 85 genes were associated with lipid metabolism with 40 genes upregulated and 45 genes downregulated (Table 1).

**Table 1.** DGEs analysis based on functional categories in wild type.

| Category | Cholesterol |  |  | Mycolic acid |  |  |
| --- | --- | --- | --- | --- | --- | --- |
|  | Total | Up | Down | Total | Up | Down |
| Lipid metabolism | 109 | 56 | 53 | 85 | 40 | 45 |
| Conserved<br>hypotheticals/Unknown | 378 | 221 | 157 | 250 | 127 | 123 |
| Cell wall and cell processes | 295 | 136 | 159 | 183 | 99 | 84 |
| Information pathways | 88 | 25 | 63 | 45 | 14 | 31 |
| Intermediary metabolism and<br>respiration | 306 | 154 | 152 | 187 | 108 | 79 |
| Regulatory proteins | 57 | 35 | 22 | 44 | 18 | 26 |
| Insertion seqs and phages | 42 | 24 | 18 | 30 | 12 | 18 |
| PE/PPE | 61 | 27 | 34 | 60 | 43 | 17 |
| Virulence, detoxification,<br>adaptation | 72 | 44 | 28 | 54 | 17 | 37 |

The number of differentially expressed genes are shown by functional categories in cholesterol and mycolic acid and DGEs are based on log2 fold change of $\geq 1$ or $\leq -1$ and adjusted p-value/false recovery rate of $\leq 0.05$ .

**Fig. 2.**
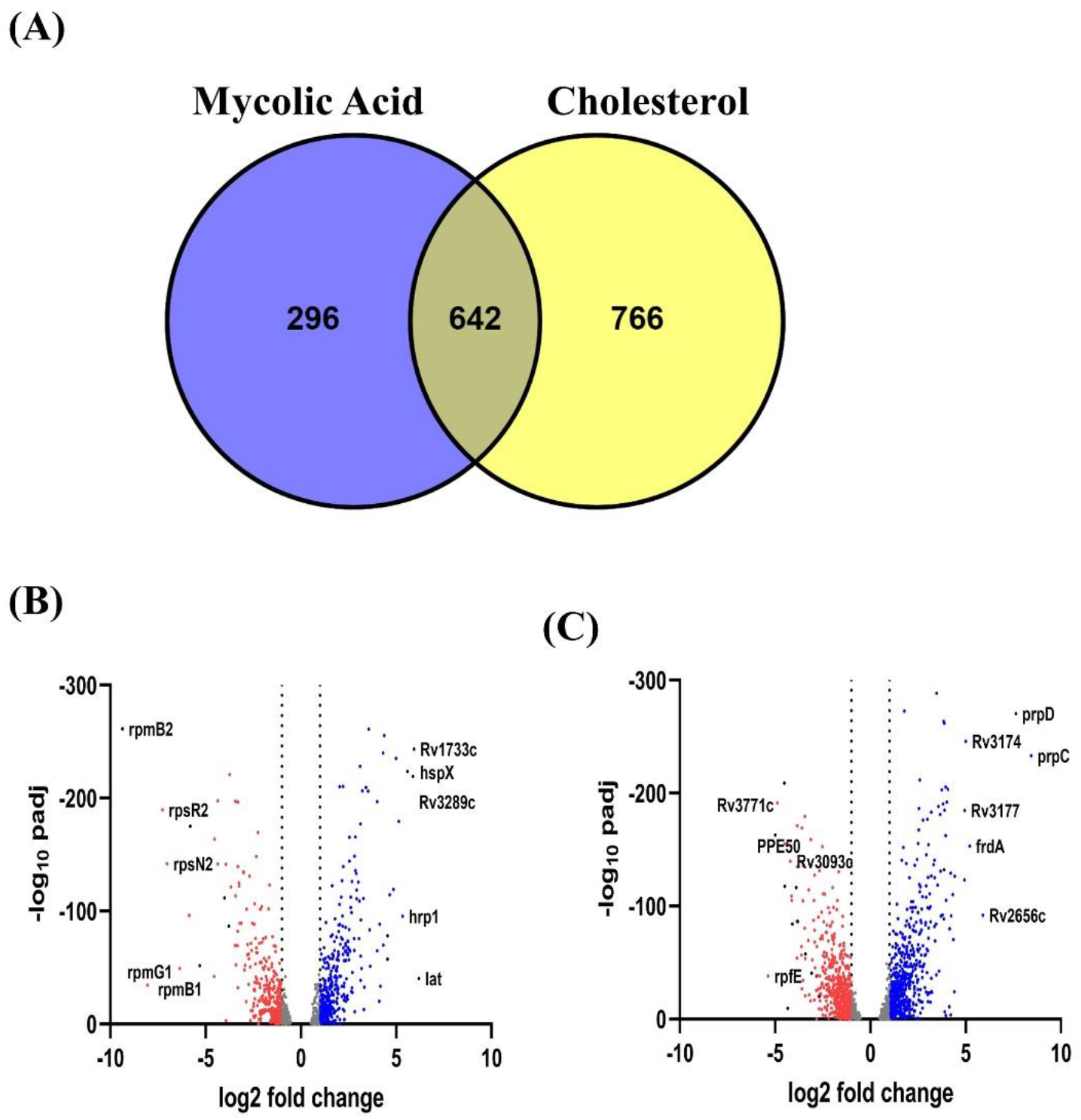
Venn diagram and volcano plot of DGEs in wild type Mtb. Venn diagram showing a total of 1408 DGEs in Cholesterol and 938 DGEs within mycolic acid condition compared to glycerol as a sole carbon source. 642 DGEs were common between these two conditions B) Volcano plot showing DGEs in cholesterol. C) Volcano plot showing DGEs in mycolic acid. In the volcano plot, X-axis represents the log of the fold change (base 2) and Y-axis represents the negative log of false discovery rate (FDR) (base 10). The red points represent downregulated transcripts (FC ≤ −1 and FDR ≤ 0.05) and the blue points represent transcripts upregulated in each condition (FC ≥ 1) and FDR ≤ 0.05).

When comparing the differentially expressed genes between the wild type and *Δmkl* strains grown under similar carbon sources, minimal differences were observed when glycerol is used as the carbon source (30 genes) followed by moderate differences in the presence of mycolic acid (216 genes) and a significantly higher number of genes differentially expressed in the presence of cholesterol (1013 genes) (Table 2). The functional categories of DGEs fell into the same categories of conserved hypothetical proteins/unknown, cell wall and cell processes, and intermediary metabolism and respiration (Table 2) as the wild type strain grown in the presence of mycolic acid or cholesterol. Among the differentially expressed genes, only 20 (13 upregulated and 7 downregulated) and 68 (31 upregulated and 37 downregulated) genes were related to lipid metabolism in the presence of mycolic acid and cholesterol, respectively. Notably, 126 DGEs were found to be common between these two conditions (cholesterol and mycolic acid) (Fig. 3).

**Table 2.** DGEs analysis based on functional categories in *Δmkl*.

| Category | Glycerol |  |  | Mycolic acid |  |  | Cholesterol |  |  |
| --- | --- | --- | --- | --- | --- | --- | --- | --- | --- |
|  | Total | Up | Down | Total | Up | Down | Total | Up | Down |
| Lipid metabolism | 3 | 1 | 2 | 20 | 13 | 7 | 68 | 31 | 37 |
| Conserved<br>hypotheticals/Unknown | 3 | 2 | 1 | 52 | 24 | 28 | 259 | 110 | 149 |
| Cell wall and cell processes | 7 | 4 | 3 | 45 | 26 | 19 | 218 | 116 | 102 |
| Information pathways | 2 | 2 | 0 | 12 | 6 | 6 | 57 | 31 | 26 |
| Intermediary metabolism<br>and respiration | 4 | 4 | 0 | 42 | 20 | 22 | 229 | 99 | 130 |
| Regulatory proteins | 5 | 3 | 2 | 8 | 2 | 6 | 38 | 11 | 27 |
| Insertion seqs and phages | 2 | 1 | 1 | 8 | 5 | 3 | 26 | 10 | 16 |
| PE/PPE | 1 | 0 | 1 | 17 | 12 | 5 | 42 | 22 | 20 |
| Virulence, detoxification,<br>adaptation | 3 | 0 | 3 | 12 | 7 | 5 | 76 | 27 | 49 |

The number of differentially expressed genes are shown by functional categories in *Amkl* mutant compared to wild type in glycerol, mycolic acid and cholesterol and DGEs are based on log2 fold change of $\geq 1$ or $\leq -1$ and adjusted p value/false recovery rate of $\leq 0.05$ .

**Fig. 3.**
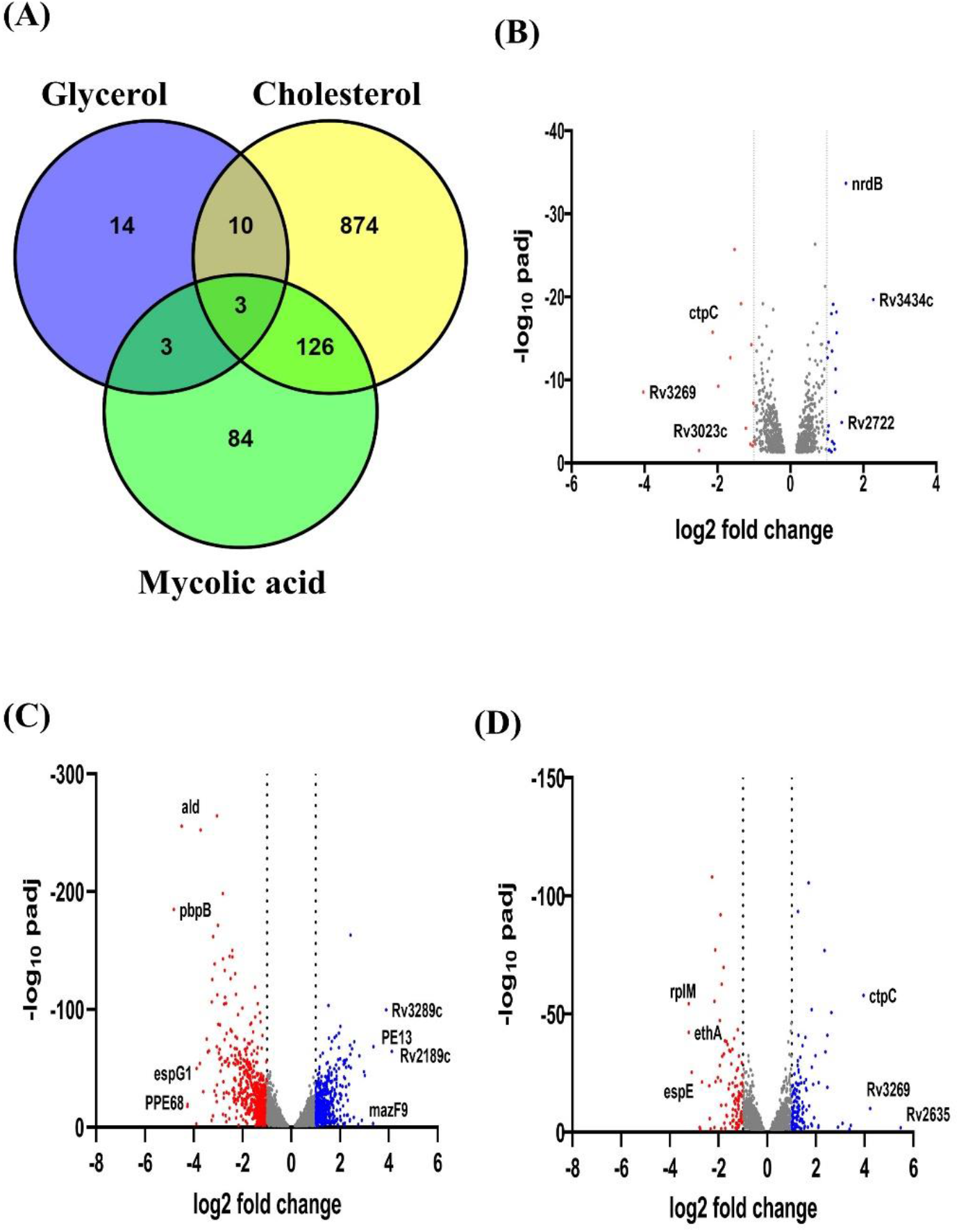
Venn diagram and volcano plot of DGEs in *Δmkl*. A) Venn diagram showing DGEs in glycerol, mycolic acid and cholesterol in *Δ*mkl when compared to wild type grown using the same carbon source. Volcano plot B) Glycerol C) Cholesterol D) Mycolic acid, where X-axis represents the log of the fold change (base 2) and Y-axis represents the negative log of false discovery rate (FDR) (base 10). The red points represent downregulated transcripts (FC ≤−1 and FDR < 0.05) and the blue points represent transcripts upregulated in each condition (FC ≥ 1 and FDR < 0.05).

### 3.2 Effect of cholesterol and mycolic acid on *mce* operon

The effects of cholesterol and mycolic acid as the only carbon sources on the expression of 42 *mce* genes (13 from *mce1* operon, 9 from *mce2* operon, 10 from *mce3* operon, and 10 from *mce4* operon) were investigated. Additionally, we also examined the influence of the genes of *mce* operon due to the deletion of *mkl* under different carbon source conditions (Fig. 4A). In the wild type strain, in the presence of mycolic acid, genes from *mce1* and *mce3* operons were upregulated while the genes from *mce2* and *mce4* operons were downregulated. Significantly upregulated genes include *fadD5* and *yrbe1A* and on the other hand, *yrbE4B* was significantly downregulated. When cholesterol was used as the carbon source, upregulation of expression was observed for the genes from *mce1, mce2*, and *mce3* operons, whereas the expression of *mce4* operon genes were downregulated. *fadD5, yrbE3A, yrbE3B, mce3A-F*, and *mam3A* were significantly upregulated. Conversely, *yrbE4A, yrbE4B, mce4C-F, mam4A*, and *mam4B* were significantly downregulated. Interestingly, *mkl* was found to be upregulated in the presence of both, cholesterol as well as mycolic acid.

**Fig. 4.**
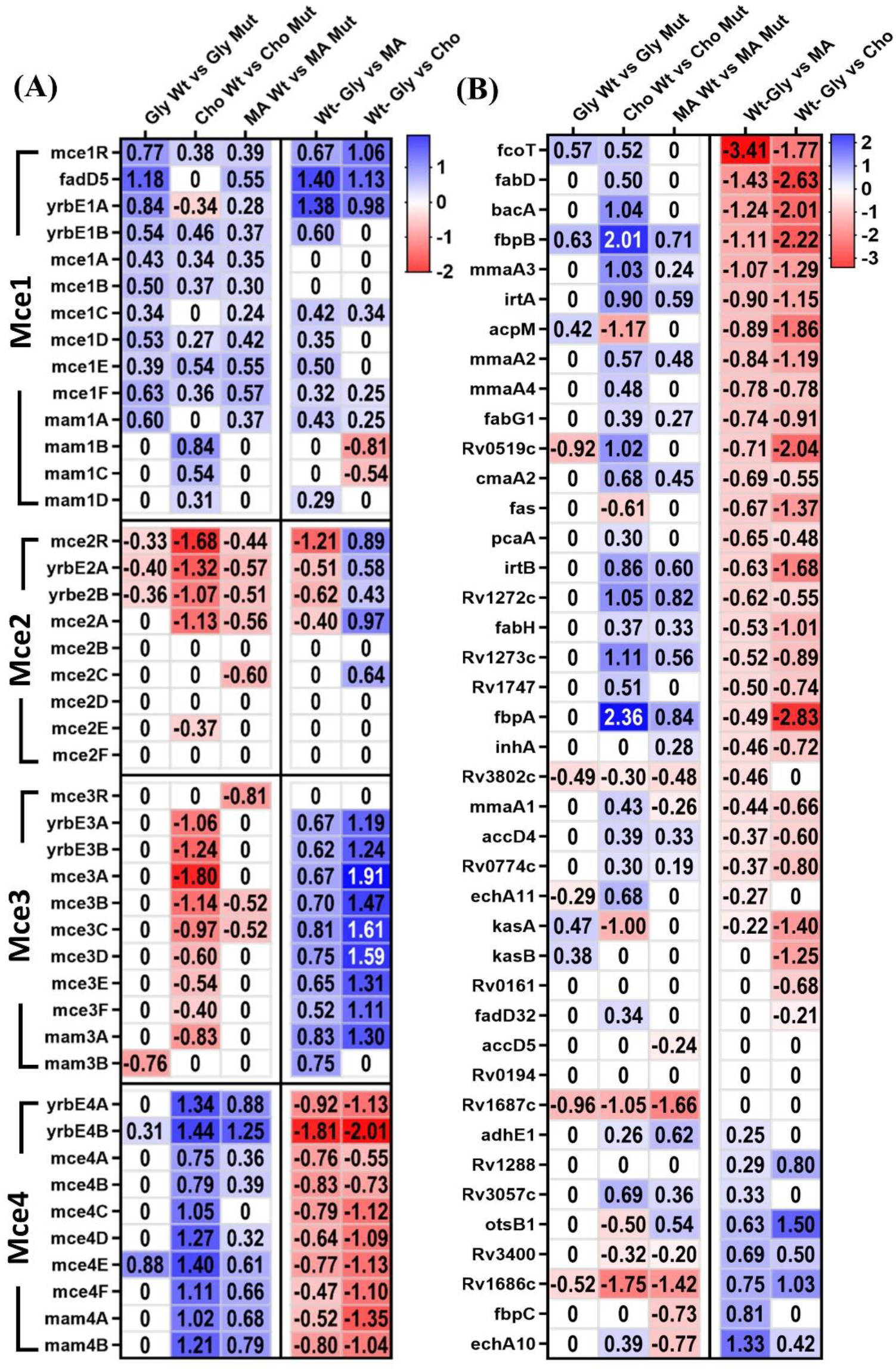
Heatmap with DGEs in mce operon genes and mycolic acid metabolizing genes. Heatmap with the DGEs with the log2-fold change A) mce operon genes and B) Mycolic acid metabolizing genes.

When comparing the expression profile of the *Δmkl* strain in glycerol, *fadD5* was upregulated in the *Δmkl* strain compared to that in the wild type strain. On a similar analysis with only mycolic acid as the sole carbon source, *yrbE4B* from the *mce4* operon exhibited a significant upregulation. Similarly, when comparing the expression profile of the *Δmkl* strain in cholesterol, upregulation of expression of genes was observed in the *mce4* operons in *Δmkl* strain, while the *mce2* and *mce3* operons experienced downregulation. It is important to highlight that all genes within the *mce4* operon, except for *mce4A* and *mce4B*, underwent significant upregulation.

### 3.3 Effects on mycolic acid biosynthesis-related genes

To assess the potential impact of the introduced carbon source on the genes related to the metabolism of mycolic acid, their expression profiles were compared. Based on prior research by Marrakchi et al., 2014, a panel of 41 genes that exhibited potential relevance to mycolic acid metabolism in Mtb were used for this analysis [5]. When using mycolic acid as the sole carbon source, only *echA10* showed a significant upregulation in the expression. In the presence of a cholesterol-rich environment, *otsB1* and *rv1686c* were upregulated. Furthermore, in *Δmkl* strain, *rv1686c* and *rv1687c* showed downregulation of expression when mycolic acid is used as a carbon source compared to the wild type strain. Similarly, when cholesterol was used as the carbon source, *Δmkl* strain showed an upregulation in the expression of *rv0519c, mmaA3, bacA, rv1272c, rv1273c, fbpB*, and *fbpA* compared to the wild type strain (Fig. 4B).

### 3.4 Effect on cholesterol metabolism-related genes

To investigate the potential impact of specific carbon sources on cholesterol metabolism, an analysis was conducted focusing on a set of 50 well-established genes associated with cholesterol metabolism excluding *mce4* operon genes [20, 26] (Fig. 5A). Upon exposure to cholesterol as a carbon source, distinct patterns of gene expression emerged. Among the genes examined, six demonstrated negligible change in expression levels, while 23 exhibited notable upregulation. Furthermore, in the presence of mycolic acid, 15 genes remained unchanged in terms of expression levels. Interestingly, six genes exhibited upregulation, while only *echA9* demonstrated downregulation. In the context of *Δmkl* compared to the wild type strain using cholesterol as the carbon source, a set of 11 genes exhibited no significant differential expression. In the presence of mycolic acid, 25 genes maintained their expression levels without significant variation. Noteworthy, *fadB3* and *fadE18* were found to be downregulated in *Δmkl*, while *fadB* was observed to be upregulated. Lastly, under conditions involving glycerol, the gene *fadE24* exhibited a downregulation in expression.

**Fig. 5.**
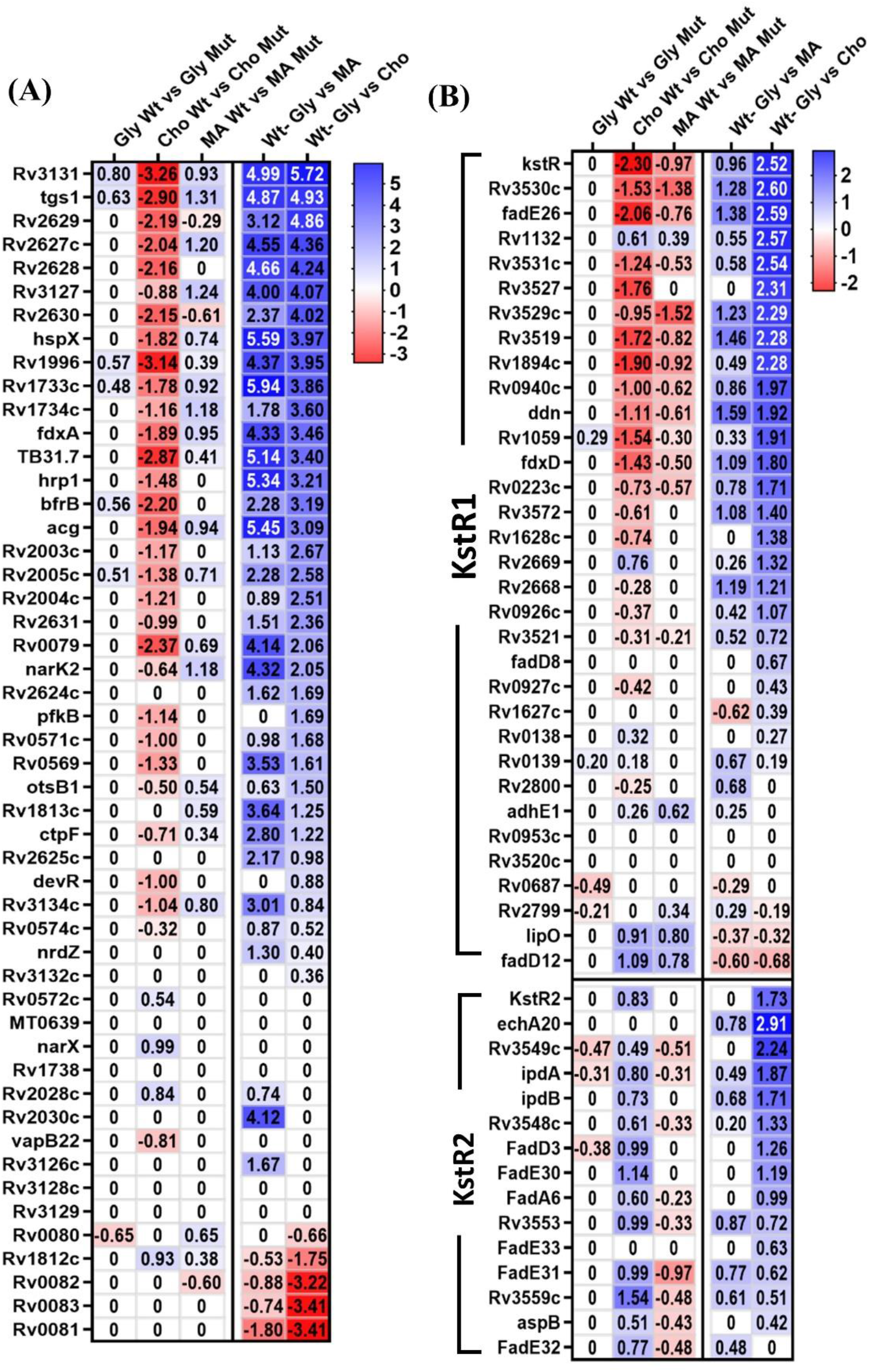
Heatmap with DGEs in cholesterol metabolizing, KstR1 and KstR2 genes. Heatmap with the DGEs with the log2-fold change A) cholesterol metabolizing genes and B) KstR1 and KstR2 regulon genes.

#### 3.4.1 kstR1 and kstR2

We conducted a comprehensive analysis of 33 genes related to *kstR1* and 15 genes related to *kstR2*, key transcriptional regulators in Mtb apart from others mentioned in cholesterol metabolism genes and *mce4* operon genes [26, 39], aiming to elucidate their differential expression profiles in response to cholesterol and mycolic acid stimuli. Among the analyzed *kstR1* genes, 19 genes exhibited upregulation in response to cholesterol. On the other hand, 8 genes showed upregulation under the influence of mycolic acid. Nonetheless, in *Δmkl*, distinct gene expression patterns were observed. *rv3529c* and *rv3530c* underwent downregulation under mycolic acid conditions. In cholesterol-rich conditions, the mutant strain exhibited upregulation of *fadD12* compared to the wild type strain. (Fig. 5B).

Within the cohort of 15 examined genes within the *kstR2* repressor, for 8 genes a trend of upregulation was noted exclusively in the presence of cholesterol. However, no significant upregulation or downregulation was noted in the presence of either mycolic acid in wild type or mutants cultured under conditions of cholesterol or mycolic acid (Fig. 5B).

### 3.5 Effect of DosR regulon

Fifty *dosR* (*devR, rv3133c*) genes associated with redox balance and recognized as key regulators of hypoxia were subjected to analysis [28, 40]. Among them, 29 genes exhibited upregulation in the presence of cholesterol. In the context of involving mycolic acid, 30 genes displayed upregulation while *rv0081* showed downregulation. Within the *Δmkl* under mycolic acid conditions, upregulation was observed in *rv1734c, narK2, rv2627c, rv3127* and *tgs1*. Interestingly, under cholesterol conditions in *Δmkl*, 24 genes exhibited downregulation in the mutant (Fig. 6A).

**Fig. 6.**
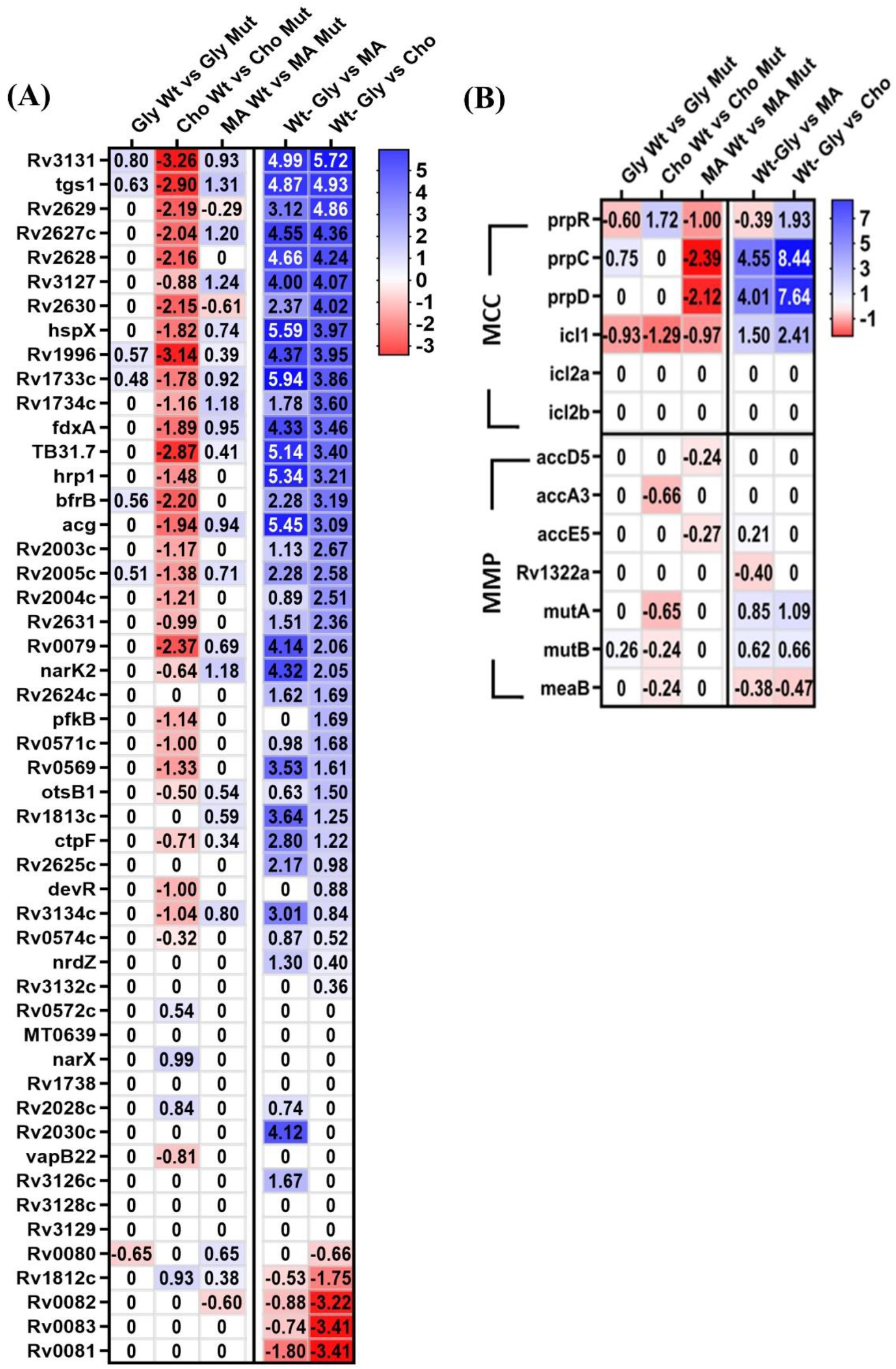
Heatmap with DGEs in DosR, MCC/MMP genes. DGEs with the log2-fold change on A) DosR regulon genes and B) MCC/MMP cycle regulon genes.

### 3.6 Effect on MCC and MMP

Propionyl-CoA, originating from cholesterol metabolism, serves as a key energy source for central metabolism through the methyl-citrate cycle (MCC) or the methyl-malonyl pathway (MMP), bypassing TCA [41]. Functional MCC is crucial for Mtb growth on cholesterol and in macrophages. Carboxylated propionyl-CoA from cholesterol could be incorporated into PDIM and SL-1 [42]. Cholesterol and mycolic acid have significantly induced the MCC cycle but not the MMP cycle. *prpC, prpD*, and *icl1* showed upregulation in response to both cholesterol and mycolic acid while *prpR* showed upregulation only in the presence of cholesterol. In the *Δmkl* strain, upregulation was observed only for *prpR* in the presence of cholesterol as the carbon source compared to the wild type strain. However, interestingly, there was significant downregulation of *prpC* and *prpD* in the presence of mycolic acid in the *Δmkl* strain compared to that in the wild type. PrpR is a transcription factor that plays a direct role in controlling the genes responsible for encoding important enzymes in the MCC cycle, namely PrpD, PrpC and Icl1 [43, 44]. The initial step of the glyoxylate cycle is mediated by Icl1, which transforms isocitrate into glyoxylate and succinate, while concurrently participating in the methylcitrate cycle to produce succinate from 2-methylisocitrate, highlighting Icl’s dual role crucial for mycobacterial fatty acid metabolism [45]. The unexplored function of the methylmalonyl pathway poses an intriguing alternative to the MCC, raising questions about its role in propionate metabolism in Mtb [46, 47]. None of the genes, except *mutA* in cholesterol conditions compared to glycerol, exhibited upregulation in the tested conditions, aligning with the findings of a prior experiment conducted by Pawelczyk et al. [26] (Fig 6B).

### 3.7 Function and pathway enrichment analysis

For a better understanding of the differentially expressed genes in each condition tested, we have used DAVID online tool. The function enrichment analysis was based on GO databases and the DGEs were significantly enriched in biological processes, molecular function and cellular component categories (Table S1).

In cholesterol, upregulated DGEs show significant enrichment in response to hypoxia (GO:0001666), cholesterol catabolic process (GO:0006707), protein secretion by the type VII secretion system (GO:0044315), cell wall (GO:0005618), oxidoreductase activity (GO:0016491), long chain-alcohol O-fatty-acyltransferase activity (GO:0047196) and response to nitric oxide (GO:0071731). The most significantly enriched pathways are associated with microbial metabolism in diverse environments (mtu01120). In mycolic acid, upregulated DGEs show significantly enriched response to hypoxia (GO:0001666), long chain-alcohol O-fatty-acyltransferase activity (GO:0047196) and response to acidic pH (GO:0010447) and the pathways include metabolic pathways (mtu01100), glycerolipid metabolism (mtu00561) and fatty acid degradation (mtu00071). We also identified 642 genes that were common between the cholesterol and mycolic acid conditions and enrichment analysis for these uniquely altered genes revealed that the top significantly enriched category was response to hypoxia (GO:0001666).

Lastly, we examined the enrichment and pathway analysis for the *Δmkl* strain grown in glycerol, mycolic acid and cholesterol. *Δmkl* strain grown in cholesterol, upregulated DGEs are significantly enriched in cell wall organization (GO:0071555) and the enriched pathways include arabinogalactan biosynthesis (mtu00572) and homologous recombination (mtu03440). *Δmkl* strain grown in cholesterol, upregulated DGEs show significant enrichment in lipopolysaccharide biosynthesis (mtu00540) pathway.

## 4. Discussion

Mtb uses a variety of physiological mechanisms and modulates both its own and the host’s lipid metabolism, in order to consume exogenous fatty acids and cholesterol [14]. However, little is understood about the metabolic changes that microbes undergo, both in terms of substrate and metabolic pathway adaptations, while responding to challenging circumstances inside the host [48].

In a comparative analysis of gene expression in Mtb H37Rv, 1408 genes exhibited differential expression in cholesterol, while 938 genes in mycolic acid. Interestingly, when comparing wild type and *Δmkl* strains in glycerol, minimal changes were observed, with only 30 DGEs. Conversely, significant alterations were observed when grown on mycolic acid, with 216 DGEs, while a remarkable shift was observed on cholesterol, with 1013 DGEs. We analyzed DGEs to focus on the *mce* operon, regulatory genes *kstR1* and *kstR2*, genes associated with cholesterol and mycolic acid metabolism, as well as genes involved in MMP and MCC.

Mtb *mce* operons may be connected to organ- and stage-specific gene functions [22, 49] and their expression may be affected by different environmental and experimental circumstances [49]. In our study, notably, we observed significant upregulation of *fadD5* from the *mce1* operon in both conditions with the media containing either cholesterol or mycolic acid. The increase in FadD5 expression related to mycolic acid implies a potential role of FadD5 in mycolic acid recycling or metabolism as also observed in earlier studies [10]. Previous studies show that the expression of *mce2* operon genes was not detected [49], which aligned to our study. In our investigation, among the *mce* operons, significant changes in expression patterns were observed mainly in *mce3* and *mce4* operons, especially in the presence of cholesterol. All of the genes of *mce3* operon were significantly upregulated in cholesterol except *mam3B. mce3* operon is transcriptionally regulated by *mce3R* but the inducer/condition for the regulation is not known [50]. The upregulation of *mce3* operon is an intriguing finding but needs further exploration to understand its effect in cholesterol metabolism/transport. In contrast, despite the strong evidence for the importance of *mce4* operon in cholesterol transport [18, 21], our research revealed that the *mce4* operon did not upregulate in the presence of cholesterol. Instead, the expression of *mce4* operon genes is downregulated. However, the observation of the downregulation of *mce4* operon genes is consistent with prior studies when cholesterol was used as the sole carbon source in Mtb [26] and *G. neofelifaecis* [51]. Prior research has also demonstrated that deleting *mce4* genes only partially hinders cholesterol transport but cholesterol presence does not significantly affect their expression pattern [26]. There can be multiple reasons for this. For example, it is possible that the transport activity of Mce4 complex, but not the amount of Mce4 complex itself, is increased in the presence of cholesterol. Also, there could be other transporters involved in the import of cholesterol or even other mechanisms of cholesterol import not yet known. Therefore, our data suggests that Mtb *mce* operon control may be more complex than expected and *mce* operon genes may be regulated differently as also reported earlier [52]. Another interesting observation is that in the *Δmkl* strain, *mce3* operon is downregulated while *mce4* operon is upregulated, a phenomenon also requiring further studies.

The lack of a direct effect on the expression levels of *mce4* operon triggered us to further analyze other known genes involved in cholesterol metabolism. According to earlier research, genes related to cholesterol metabolism were shown to be upregulated for survival inside macrophages [53, 54]. In the presence of cholesterol, 23 out of 50 genes exhibited upregulation which also largely matched with the previous study [26]. However, intriguingly, in the *Δmkl* strain, several of these genes were downregulated when compared to the wild type expression levels. The cholesterol catabolic cluster is thought to be comprised of four *fadD* genes: *fadD3, fadD17, fadD18*, and *fadD19*, among the *fadD*s associated with lipid synthesis and degradation [53]. In our present study, indeed *fadD3, fadD17* and *fadD19* showed upregulation when exposed to cholesterol. Notably, only *fadD17* showed upregulation in the presence of mycolic acid. The upregulation of these genes when exposed to cholesterol is in agreement with previous studies conducted by Nesbitt et al. [20] and Pawelczyk et al. [26]. In general, our observation was that the effect of mycolic acid in the media followed a similar pattern as that of cholesterol for the expression of cholesterol metabolism related genes but to a lesser extent. This could be a general response of the mycobacteria to lipids as carbon and energy source.

Upon examining the genes associated with mycolic acid metabolism, very minor change was observed in the presence of mycolic acid itself. We speculate that free mycolic acid does not change the expression of enzymes necessary for de-novo mycolic acid biosynthesis. The precise mechanisms of mycolic acid recycling in Mtb are not fully understood. In the *Δmkl* knockout strain, upregulation of few genes were observed exclusively in the presence of cholesterol as the carbon source compared to the wild type strain. *bacA, rv1272c*, and *rv1273c*, all belonging to the ABC transporter family [55-57], exhibit upregulation in the *Δmkl* knockout strain compared to the wild type, indicating the presence of functional redundancy. This also suggests the potential existence of other transporters compensating for the effects of *mkl* in the mutant.

During the breakdown of fatty acids or cholesterol, changes in metabolic pathways involving propionyl-CoA and pyruvate led to the necessity of inducing transcription of MCC enzymes for growth on this substrate [41, 58]. Our study shows that Mtb’s consumption of cholesterol and mycolic acid induces MCC and glyoxylate shunt. MCC gene upregulation has also been reported in previous studies [26, 59]. Interestingly, the mycolic acid metabolism induces upregulation of MCC cycle-related genes.

The dormancy regulon DosR system is a vital regulator of hypoxia in Mtb and also involved in maintaining redox balance, protecting Mtb from oxidative stress [40, 60]. Interestingly, the activation of DosR is not solely dependent on hypoxia, the presence of cholesterol in the Mtb growth environment can strongly induce the dormancy related response [28]. 36 of the DosR-regulated genes were reported to be upregulated when cholesterol was used as a carbon source [26]. In our present study, cholesterol upregulated 29 genes and mycolic acid upregulated 30 genes. The upregulation of the genes related to dormancy in both cholesterol and mycolic acid states the importance of those fatty acids for the persistence of Mtb and how Mtb metabolizes those available carbon sources. Due to a limited understanding of Mtbs metabolic activity during latency, developing drugs for dormant bacilli is challenging [28]. We speculate that Mtb might similarly metabolize cholesterol and mycolic acid under stress conditions. Cholesterol metabolism becomes important during the persistence stage of infection rather than the growth phase [20]. It might also be true for recycling mycolic acid.

## 5. Concluding remarks

In contrast to prior research, which often focused on mixed carbon sources, this study uniquely emphasizes specific carbon substrates especially on cholesterol and mycolic acid, shedding light on Mtb’s adaptive strategies and regulatory mechanisms. The identification of genes affected by the presence of cholesterol and mycolic acid and their potential crosstalk provides a clear view of Mtb’s survival strategies within the host. Mkl could have a broader effect on lipid transport and metabolism than ever thought. Both cholesterol and mycolic acid (mycolic acid could be better) containing media could be used as an *in vitro* model of dormant Mtb for drug discovery and drug testing providing a novel perspective.

## Supporting information

RNA seq data

## Abbreviations

Mtb: Mycobacterium tuberculosis
MCE: mammalian cell entry
RNAseq: RNA sequencing
DEG: differentially expressed gene

## Data availability statement

All data generated in this study are included in this article.

## Supplementary Material

The following is the Supplementary data to this article: Table S1: Gene Ontology (GO) and KEGG enrichment analyses of the RNAseq data (DAVID Bioinformatics Resources), RNAseq result analysis.

## CRediT authorship contribution statement

**Mohammad Asadur Rahman**: Writing–original draft, Methodology, Formal analysis, Investigation, Visualization. **Lee W. Riley**: Conceptualization, Supervision, Funding acquisition. **Valerio Iizzi**: Formal analysis, Data curation, Validation, Writing, Review & editing. **Rajaram Venkatesan**: Writing, Review & editing, Supervision, Funding acquisition, Conceptualization.

## Acknowledgments

We acknowledge the support from the QB3-Berkeley Genomics facility at the University of California, Berkeley for RNAseq data collection.

## Ethical statement

Not applicable.

## Consent for publication

Not applicable.

