## Supplementary data for "Mycolic acid and cholesterol induced transcriptomic adaptations in wild type and Mce complex ATPase subunit deleted strain of *M. tuberculosis* H37Rv"

**Table S1.** Gene Ontology (GO) and KEGG enrichment analyses of the RNAseq data (DAVID Bioinformatics Resources).

| Experimental condition | ID and Term | Category | No of genes | P Value |
|---|---|---|---|---|
| Cho upregulated | GO:0001666~response to hypoxia | Biological process | 27 | 7,72E-09 |
| | GO:0006707~cholesterol catabolic process | Biological process | 17 | 0,00000686 |
| | GO:0044315~protein secretion by the type VII secretion system | Biological process | 9 | 0,0000135 |
| | GO:0005618~cell wall | Cellular component | 134 | 0,001355789 |
| | GO:0016491~oxidoreductase activity | Molecular function | 39 | 0,001940596 |
| | GO:0047196~long chain-alcohol O-fatty-acyltransferase activity | Molecular function | 9 | 0,002128348 |
| | GO:0071731~response to nitric oxide | Biological process | 9 | 0,002571333 |
| | mtu01120:Microbial metabolism in diverse environments | KEGG pathway | 52 | 4,16958E-06 |
| | mtu00561:Glycerolipid metabolism | KEGG pathway | 13 | 0,000156815 |
| | mtu00620:Pyruvate metabolism | KEGG pathway | 17 | 0,000194081 |
| | mtu00984:Steroid degradation | KEGG pathway | 8 | 0,000600832 |
| | mtu02020:Two-component system | KEGG pathway | 17 | 0,002659671 |
| Cho downregulated | GO:0071555~cell wall organization | Biological process | 26 | 7,72913E-05 |
| | GO:0004315~3-oxoacyl-[acyl-carrier-protein] synthase activity | Molecular function | 11 | 0,000807363 |
| | GO:0097040~phthiocerol biosynthetic process | Biological process | 6 | 0,001298085 |
| | mtu03440: Homologous recombination | KEGG pathway | 14 | 7,04623E-05 |
| | mtu00572: Arabinogalactan biosynthesis - Mycobacterium | KEGG pathway | 9 | 0,002658628 |
| MA upregulated | GO:0001666~response to hypoxia | Biological process | 23 | 5,88936E-09 |
| | GO:0047196~long chain-alcohol O-fatty-acyltransferase activity | Molecular function | 9 | 0,000160413 |
| | GO:0010447~response to acidic pH | Biological process | 8 | 0,0001929 |
| | GO:0071731~response to nitric oxide | Biological process | 9 | 0,000218457 |
| | GO:0019432~triglyceride biosynthetic process | Biological process | 9 | 0,000355611 |
| | GO:0009986~cell surface | Cellular component | 14 | 0,000452985 |
| | GO:0004144~diacylglycerol O-acyltransferase activity | Molecular function | 9 | 0,001506802 |
| | GO:0006071~glycerol metabolic process | Biological process | 9 | 0,001740378 |
| | GO:0044315~protein secretion by the type VII secretion system | Biological process | 6 | 0,002659514 |
| | mtu01100:Metabolic pathways | KEGG pathway | 82 | 4,42899E-06 |
| | mtu00561:Glycerolipid metabolism | KEGG pathway | 10 | 0,00051487 |
| | mtu00071:Fatty acid degradation | KEGG pathway | 13 | 0,000674291 |
| | mtu00310:Lysine degradation | KEGG pathway | 11 | 0,001651088 |
| | mtu00380:Tryptophan metabolism | KEGG pathway | 11 | 0,002976779 |
| MA downregulated | GO:0051607~defense response to virus | Biological process | 9 | 3,44469E-07 |
| | GO:0071770~DIM/DIP cell wall layer assembly | Biological process | 13 | 6,13714E-05 |
| | GO:0031177~phosphopantetheine binding | Molecular function | 11 | 0,000220465 |
| | GO:0004519~endonuclease activity | Molecular function | 9 | 0,000573771 |
| | GO:0097041~phenolic phthiocerol biosynthetic process | Biological process | 5 | 0,000962317 |
| | GO:0097040~phthiocerol biosynthetic process | Biological process | 5 | 0,002611595 |

| | | | | |
|---|---|---|---|---|
| | GO:0006633~fatty acid biosynthetic process | Biological process | 16 | 0,002819327 |
| Cho Mut Upregulated | GO:0071555~cell wall organization | Biological process | 19 | 0,000167488 |
| | mtu00572:Arabinogalactan biosynthesis - Mycobacterium | KEGG pathway | 9 | 0,000115948 |
| | mtu03440:Homologous recombination | KEGG pathway | 10 | 0,001047358 |
| Cho Mut downregulated | GO:0005618~cell wall | Cellular component | 132 | 1,60538E-07 |
| | GO:0001666~response to hypoxia | Biological process | 21 | 3,43594E-06 |
| | GO:0044315~protein secretion by the type VII secretion system | Biological process | 7 | 0,000682988 |
| | GO:0045893~positive regulation of transcription, DNA-templated | Biological process | 8 | 0,001675081 |
| | mtu00620:Pyruvate metabolism | KEGG pathway | 15 | 0,000878482 |
| MA Mut upregulated | mtu00540:Lipopolysaccharide biosynthesis | KEGG pathway | 3 | 0,001358898 |
| MA Mut downregulated | GO:0010033~response to organic substance | Biological process | 5 | 2,87521E-05 |
